# Quantifying Molecular Changes in the Preeclamptic Rat Placenta with Targeted Contrast-Enhanced Ultrasound Imaging

**DOI:** 10.1101/2024.12.09.627612

**Authors:** Lili Shi, Allan K. N. Alencar, Kenneth F. Swan, Dylan J. Lawrence, Gabriella Pridjian, Carolyn L. Bayer

## Abstract

**Purpose:** Abnormal placental remodeling is linked to various pregnancy-related diseases, including preeclampsia (PE). This study applies a bicompartmental (BCM) model to quantify molecular expression changes in the placenta, indicative of abnormal placental remodeling, and evaluates the effectiveness of targeted contrast-enhanced ultrasound (T-CEUS) in detecting the abnormal placental vasculature. The BCM model provides high temporal resolution and differentiation of anatomical artery structures within the placenta by analyzing the distribution of contrast agents.

**Methods:** A targeted contrast agent (TCA) composed of gas-filled microbubbles (MB), with a surface-conjugated peptide to target α_ν_β_3_ integrin, a biomarker for angiogenesis, was used for quantifying placental vascular development. CEUS images were acquired from timed pregnant Sprague Dawley rats with experimentally-induced reduced uterine perfusion pressure (RUPP) placental insufficiency. On gestational day (GD) 18 of a 21-day gestation, CEUS images were acquired from both Normal pregnant (NP; n=6) and RUPP (n=6) dams after injection of the TCA. The BCM model was used to estimate the binding dynamics of the TCA, providing a parametric map of the binding constant (*K_b_*) of the placenta.

**Results:** The RUPP group showed a significant reduction in the value of *K_b_ c*ompared to the NP group (*p* < 0.05). A histogram of the placental *K_b_ w*as compared to alternative analyses (differential target enhancement, dTE and late enhancement, LE) to demonstrate that it can differentiate between anatomical artery structures with a higher contrast-to-background ratio.

**Conclusions:** The BCM method differentiates molecular changes associated with the abnormal placental development associated with PE. It also reveals more intricate internal anatomical structures of the placenta in comparison to dTE and LE, suggesting that the BCM could enhance early detection and monitoring of PE.

## Introduction

Preeclampsia (PE) is a complex condition that contributes significantly to maternal and neonatal morbidity and mortality, affecting 6-8% of pregnancies and linked to over 70,000 deaths worldwide [1]. Symptoms of PE include high blood pressure and proteinuria, typically emerging during the second half of pregnancy [2]. The placenta is a vital organ with a complex vascular structure that supports fetal development by ensuring efficient gas and nutrient exchange [3]. Blood from the maternal uterine arteries enters the placenta through spiral arteries, filling the intervillous space where exchange between maternal and fetal blood occurs [4]. In normal development, the spiral arteries undergo significant remodeling, losing smooth muscle cells and elastic tissue, which causes them to dilate and thus increase maternal blood flow to the placenta [5]. This low-pressure, low-velocity environment allows blood to flow freely over the placental villi, facilitating the transfer of oxygen and nutrients [6, 7]. However, when placental vascular development and remodeling are inadequate, it can lead to poor placental development and restricted blood flow [6, 8–10]. These abnormalities are linked to PE, a condition with limited treatment options, where delivery of the placenta remains the only definitive solution [11]. Therefore, an in-depth analysis of placental vessel structure and function is needed to better understand the pathophysiology of PE.

Targeted contrast-enhanced ultrasound (T-CEUS) imaging uses gas-filled echogenic microbubbles (MBs) as contrast agents, which are directed to specific molecular markers by attaching suitable ligands to the MB surfaces [12, 13]. Upon intravenous administration, these functionalized MBs bind to tissues expressing the target molecular markers, resulting in a localized increase in the ultrasound imaging signal [14–16]. Due to their size, typically several micrometers, contrast MBs largely remain within the vascular system, making T-CEUS highly effective for noninvasively detecting and monitoring biological processes occurring in vascular endothelial cells, such as those involved in placental angiogenesis [14]. Achieving high-contrast images requires differentiation between MBs that are bound to molecular biomarkers (“adherent” MBs) and those that remain unbound (“free” MBs). Two specialized semi-quantitative methods have been developed: differential targeted enhancement (dTE) and late enhancement (LE) [17–19]. The dTE method enhances contrast signals from both adherent and circulating MBs after the targeted contrast agent (TCA) injection by applying a destructive ultrasound pulse to distinguish between bound and freely circulating MBs [20]. This approach has drawbacks, including potential damage to blood vessels and tissues due to the destructive pulse [20]. The LE method assumes that free circulating MBs are absent 10 minutes post-injection, indicating specific binding of targeted MBs [21]. While both dTE and LE provide qualitative insights, they are less suitable for quantitative analysis due to the extended data collection period, during which some MBs may degrade [21]. Additionally, residual free MBs may still affect the CEUS signal [13], and destructive pulses may cause unexpected biological effects on the rat [22, 23]. In light of these challenges, this study aims to introduce and validate a bicompartmental (BCM) model for analyzing the kinetics of a TCA composed of gas-filled MBs engineered to target α_ν_β_3_ integrin, a biomarker for angiogenesis and cell adhesion [24, 25], during placental perfusion. By fitting each pixel’s time-intensity curve (TIC) into a mathematical BCM model, we capture the concentration dynamics of ultrasound contrast agents over a short duration, avoiding long imaging durations and burst pulses. This model uses first-pass binding (FPB) assumptions to provide a detailed mapping of angiogenesis and MB binding parameters, including the binding constant (*K_b_*) for each pixel [20, 26]. We demonstrate that the first-pass binding BCM model offers a more precise and comprehensive analysis of vascular changes in placental tissue, ultimately enhancing the diagnosis and monitoring of PE.

## Materials and Methods

An expanded Materials and Methods section is available in the Supplemental Material.

### In Vivo Contrast-Enhanced Ultrasound Imaging

All animal studies were approved by Tulane University’s Institutional Animal Care and Use Committee and were conducted following the guidelines outlined in the NIH Guide for the Care and Use of Laboratory Animals. Timed-pregnant Sprague Dawley rats (8 weeks old, 250-350 g) were obtained from a commercial vendor (Charles River Laboratories, Boston, MA). On GD14, rats were divided into normal pregnant (NP; n = 6) and RUPP (n = 6) groups. The RUPP procedure was performed under anesthesia, with 2-3% isoflurane, and involved placing silver clips on the lower abdominal aorta and branches of the ovarian arteries to induce placental ischemia as previously published [27–29].

Vevo Targeted-Ready Micro Markers (Fujifilm, VisualSonics) were used as contrast agents. These microbubbles (MBs) can be conjugated with peptides to target specific endothelial cell biomolecules. Following the manufacturer’s protocol, the MBs were reconstituted in saline. Then, 20 µg of biotin-Arginine-Glycine-Aspartate-OH TFA salt (RGD) peptide, diluted in saline, was injected into the MB vial. The resulting TCA were designed to bind to α_ν_β_3_ integrin [30].

On GD18, the rats were anesthetized with 3% isoflurane. Depilatory cream was applied to remove hair from the abdominal area. Heart rate, respiration rate and body temperature were maintained throughout the imaging session. Ultrasound imaging data were acquired using the Vevo 2100 ultrasound system (FUJIFILM VisualSonics, Toronto, Canada) with an LZ250 linear array transducer (central frequency: 20 MHz; broadband frequency: 13-24 MHz; axial resolution: 75 µm; lateral resolution: 165 µm; field of view: 23 mm × 30 mm). B-mode ultrasound was used to locate one placenta for CEUS imaging (Figure 1a). The selection criteria for imaging were a placenta with well-defined edges on the B-mode ultrasound images as well as a location low in the abdominal cavity to minimize motion due to respiration. The midline of the placenta was determined using the origin of the umbilical cord, visualized using color Doppler ultrasound. To maximize the achievable frame rate, the width and depth of the CEUS imaging window were minimized.

**Figure 1.**
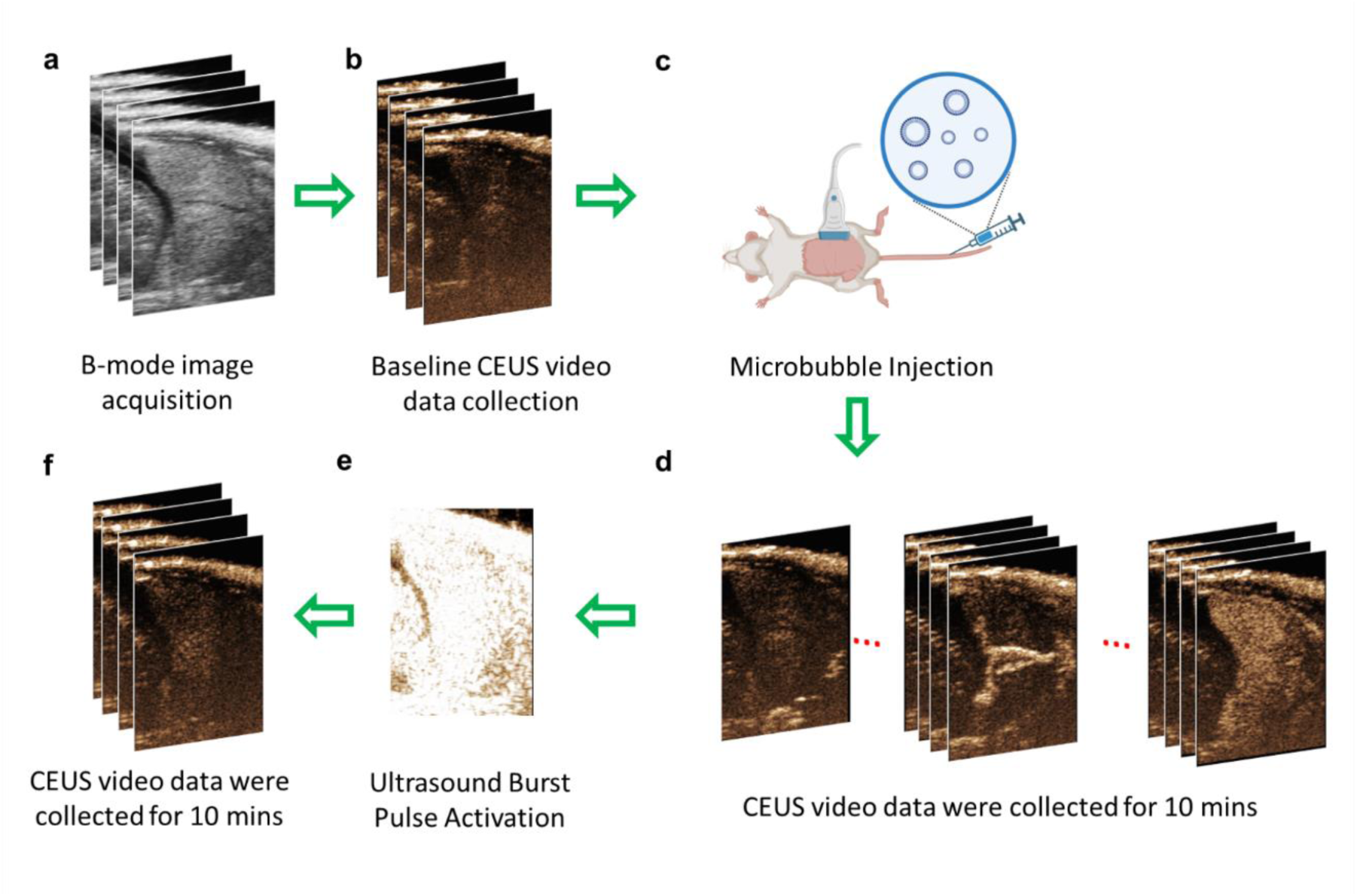
Schematic flow chart illustrating the CEUS imaging procedure. **(a)** On GD18, a B-mode image of the placenta was collected. **(b)** A baseline contrast image was collected before the injection. **(c)** After administering the contrast agent, images were collected for 10 minutes**. (d)** The microbubbles (MBs) follow the blood flow, passing through the spiral artery and central canal artery, and eventually entering the slower-flow labyrinth of the placenta. **(e)** At t=10 minutes, the ‘Burst’ mode was activated, **(f)** and another 10-minute video was continuously collected.

An experimental flow chart is depicted in Figure 1. A full video loop of B-mode (Figure 1a) and nonlinear contrast images (Figure 1b) were acquired prior to MB injection to obtain a baseline measurement of background tissue structures. A 200 µL bolus of the TCA was manually administered through a tail vein catheter (22G ×1ʺ, I.D.: 0.67mm) followed by a 100 µL saline flush while simultaneously recording nonlinear CEUS images. CEUS video loops were saved continuously over a 10 min period after injection (Figure 1d). At t=10 min, a destructive ultrasound pulse was applied (Figure 1e). Contrast data was collected for an additional 10 minutes (Figure 1f).

### Bicompartmental Model Development

The bicompartmental model (BCM) assumes that the total concentration of the contrast agents in the blood can be divided into two compartments: attached and freely circulating microbubbles [26]. The total concentration of the contrast agent can be expressed as:

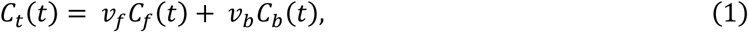

Where *C_t_*(*t*) is the total concentration of the contrast agent inside the vessel, *C_f_*(*t*) represents the concentration of free MBs, and *C_b_*(*t*) is the concentration of bound microbubbles. *v_f_* and *v_b_* correspond to the fractional volumes of free and bound MBs, respectively.

In the blood vessel, free microbubble (MB) movement is assumed to be faster than the binding kinetics of adhering MBs. Additionally, the movement of the free MBs in the longitudinal direction is influenced by the drag exerted by the flowing blood. Simultaneously, the diffusion of microbubbles is attributed to Brownian motion. The diffusion process of free microbubbles can be described by the modified local density random walk (mLDRW) model [31–33], as follows:

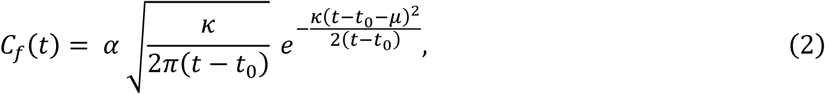

Where *μ*represents the average time taken by the free MBs to travel from the injection site to the detection site, α is the cumulative time integral of *C_f_*(*t*), *t*_0_ is the theoretical time at which the MBs are introduced into the body, and *κ* is the dispersion parameter, defined as the squared convection velocity divided by the dispersive forces.

For the bound TCA, it is assumed that all MBs will attach to the specific target molecule expressed in endothelial cells (α_ν_β_3_ integrin) and that unbinding is insignificant during the initial travel of the TCA. The binding kinetics can be represented as:

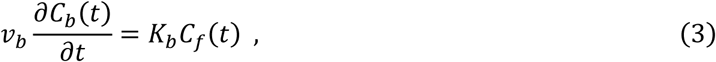

Where *K_b_* is the binding constant of bound MBs. By solving equation (3) for t≥0 and given the initial conditions *C_b_*(0) = *C_f_*(0) = 0, equation (3) can be expressed as:

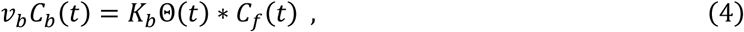

In this equation, * denotes the convolution integral, and Θ(*t*) represents the Heaviside unit-step function. By applying the adiabatic approximation [34], where the dispersion parameter κis significantly larger than *K_b_*, equation (2) is incorporated into equations (4) and (1). The entire first-pass binding BCM model can be expressed as [26]:

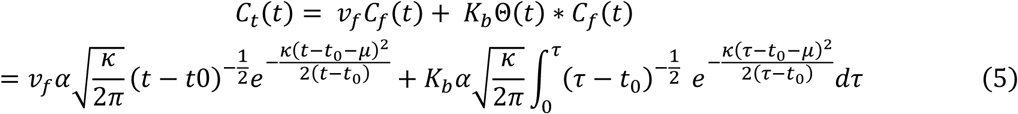

### Model Quantification

The CEUS video data from the Vevo 2100 system were converted into a MATLAB dataset. The placenta area was outlined as the region of interest (ROI) based on the B-mode image. As semi-quantitative methods, the dTE and LE approaches were used to compare quantification of MB binding. Tthe LE method assumes that the contrast in images obtained approximately 10 minutes post-injection is predominantly due to attached MBs in the placenta. A contrast image acquired 40s prior to the burst at 10 mins (at t=t_burst_-40s) was used for the LE calculation, as shown in Figure 2a.

**Figure 2.**
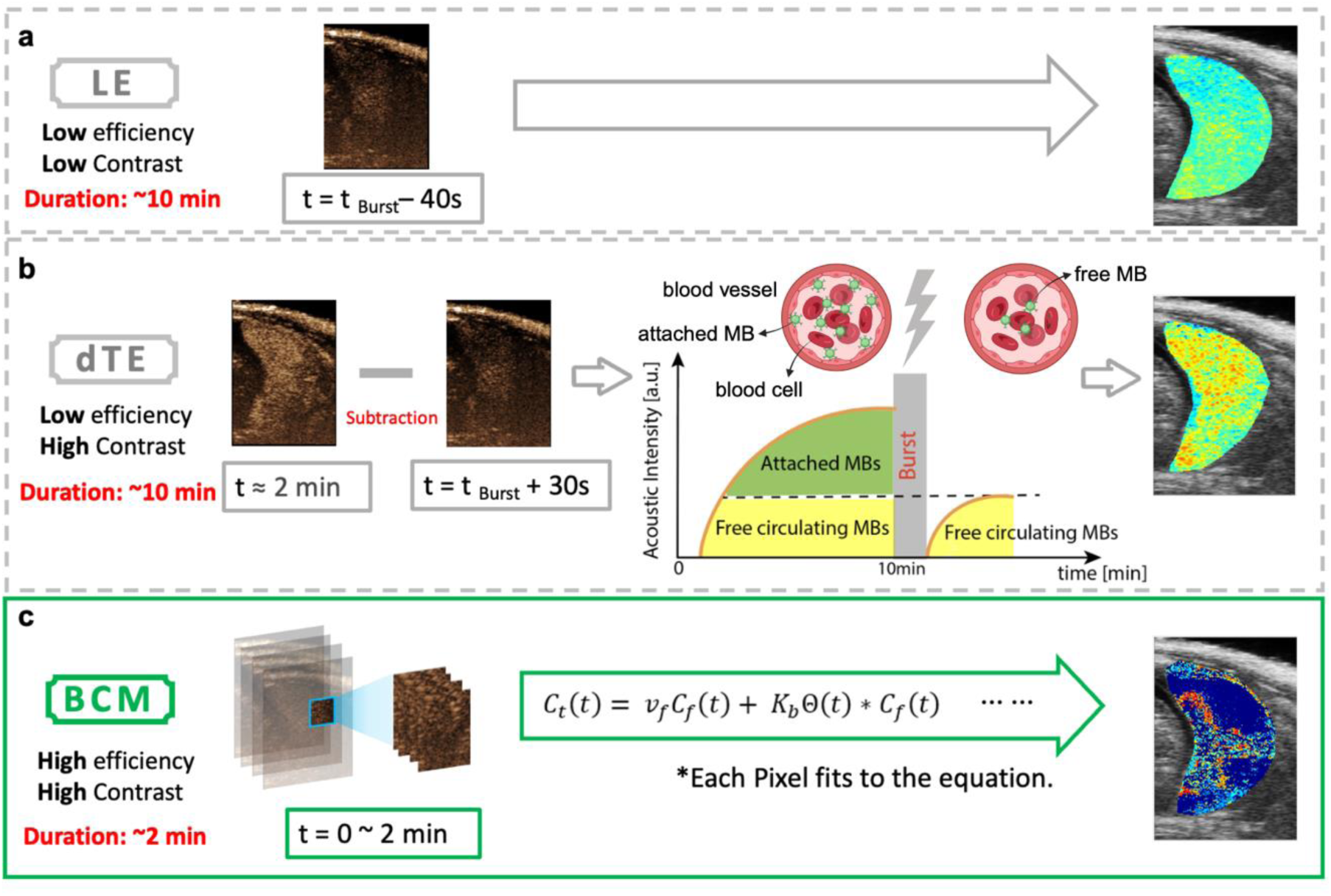
Quantification methods for binding contrast agent. **(a)** LE method: Acoustic intensity at t = t _Burst_ – 40s represents attached contrast agent value. **(b)** dTE method: Schematic showing changes before (t ≈ 2 min) and after Burst (t = t _Burst_ + 30s). The schematic illustrates the dTE method to quantify the attached contrast agent within the placenta. After tail vein injection, MBs adhere to the α_ν_β_3_ integrin on the endothelial cells. Ten minutes later, a destructive ultrasound pulse is applied to destroy the adherent MBs, and 1 minute later, the free-circulating MBs are replenished. **(c)** BCM method: data from t = 0 ∼ 2 min are fitted into the complete equation for each pixel to calculate the binding constant of the attached MBs.

The dTE method compares contrast image data at two time points: 30 seconds after a destructive ultrasound pulse (t=t_burst_+30s) and about 2 minutes post-injection (t ≈ 2min). Consistent with prior literature [20], we empirically chose t ≈ 2 mins since at this time CEUS was maximized and provided highest contrast for the placenta. For comparison, a dTEc (conventional dTE), using image frames acquired immediately pre- and post- burst is provided in t Supplementary Figure 5. The difference in acoustic intensity between these two time points represents the bound contrast agent dTE value, as shown in Figure 2b.

In contrast to the dTE and LE methods, which select a single time point, the BCM fits each pixel’s time-intensity curve (TIC) over time. A TIC was generated for each pixel of the placenta. The peak value of the linear TIC in each pixel was identified, and the curve was segmented from t_0_ to the time to peak value (t=t_0_∼2min, where t_0_ is defined as the start of contrast agent wash-in).

The segmented TICs were initially fit using the non-linear least square curve fitting method within the modified Local Density Random Walk (mLDRW) model [20]. The mLDRW fitting results provided preliminary estimates of the unknown parameters before fitting the BCM model. These segmented TIC curves were then fit to the BCM model using the non-linear least square curve fitting method. This step calculated the targeted MB binding constant (*K_b_*), with a range of (0.005; 15) min ^-1^ [26] for each pixel. Finally, a parameter map was created to visualize the distribution and intensity of MB binding across the placenta, providing a comprehensive overview of MB dynamics in the targeted area, as shown in Figure 2c.

### Molecular Expression Validation

Following euthanasia on GD18, placentas were collected, weighed, and sectioned, with one portion stored at −80 °C for subsequent real-time quantitative polymerase chain reaction (RT-qPCR) analysis to quantify mRNA expression levels of α_ν_β_3_ integrin and β-actin, while the other portion was processed for immunohistochemistry (IHC).

IHC staining was performed using the Horseradish Peroxidase method (HRP). Placental sections were incubated with mouse polyclonal α_ν_β_3_ integrin (1:200 dilution, Bioss antibodies, bs-1310R) and secondary HRP-polymer (Rabbit-On-Rodent HRP-polymer, Biocare Medical, Pacheco, CA). Slides were then incubated with 3,3′-diaminobenzidine (DAB) for 5-min and rinsed in a running water bath, and counterstained with hematoxylin. Slides were scanned using a Zeiss Axio Scan.Z1 Slide Scanner (Carl Zeiss AG, Oberkochen, Germany) and images were exported to Fiji ImageJ analysis software for processing. The expression of α_ν_β_3_ integrin was quantified as the percent area fraction of DAB staining compared to the total area of the placental section.

### Statistical Analysis

Sample sizes were determined from an a priori power analysis performed in G*Power Software (Heinrich-Heine-Universität, Dusseldorf, Germany). Statistical analysis was performed in GraphPad Prism 10.0.2 (GraphPad Software, San Diego, CA). Differences between the *K_b_*, dTE, and LE methods were compared using a one-way ANOVA with Tukey’s post-hoc test. For each method, an unpaired two sided t-test was used to compare the differences between the NP and RUPP groups. Pearson’s correlation analysis (r) assessed the linear relationships between the methods. A *p*-value < 0.05 was considered statistically significant.

## Results

### Quantitative Analysis of Targeted Contrast Agent (TCA)

CEUS data were analyzed in MATLAB to quantify binding and generate parametric maps of the bound MBs (Figure 3). Using the least square curve fitting method, the TICs of each pixel were fit to the first-pass binding BCM model to obtain the parametric maps of *K_b_*. RUPP placentas (Figure 3b) showed significantly fewer attached MBs compared to NP (Figure 3a), especially in the labyrinth zone and central canal artery. The semi-quantitative methods, dTE (Figure 3d, 3e) and LE (Figure 3g, 3h), confirmed higher values in NP placentas, consistent with BCM results (Figures 3a and 3b). Parametric maps were overlaid on the B-mode images, with histogram data presented as means ± SEM (Figure 3c, 3f, 3i). Due to the high attenuation of the ultrasound signal by the contrast microbubbles, in some acquisitions the contrast signal in the bottom half of the placenta remained below the noise threshold at the 2 min post-injection acquisition time; therefore, for all data in Figure 3c, the top half of the placenta was used to calculate the mean *K_b_*. Pixels whose signal remained at the pre-injection background signal level within the 2 min imaging time frame, indicating no microbubble perfusion, were also excluded from the calculation of the mean *K_b_* in Figure 3c. For Figure 3f and i, the B-mode image was used to identify the placental ROI, and the entire ROI was averaged.

**Figure 3.**
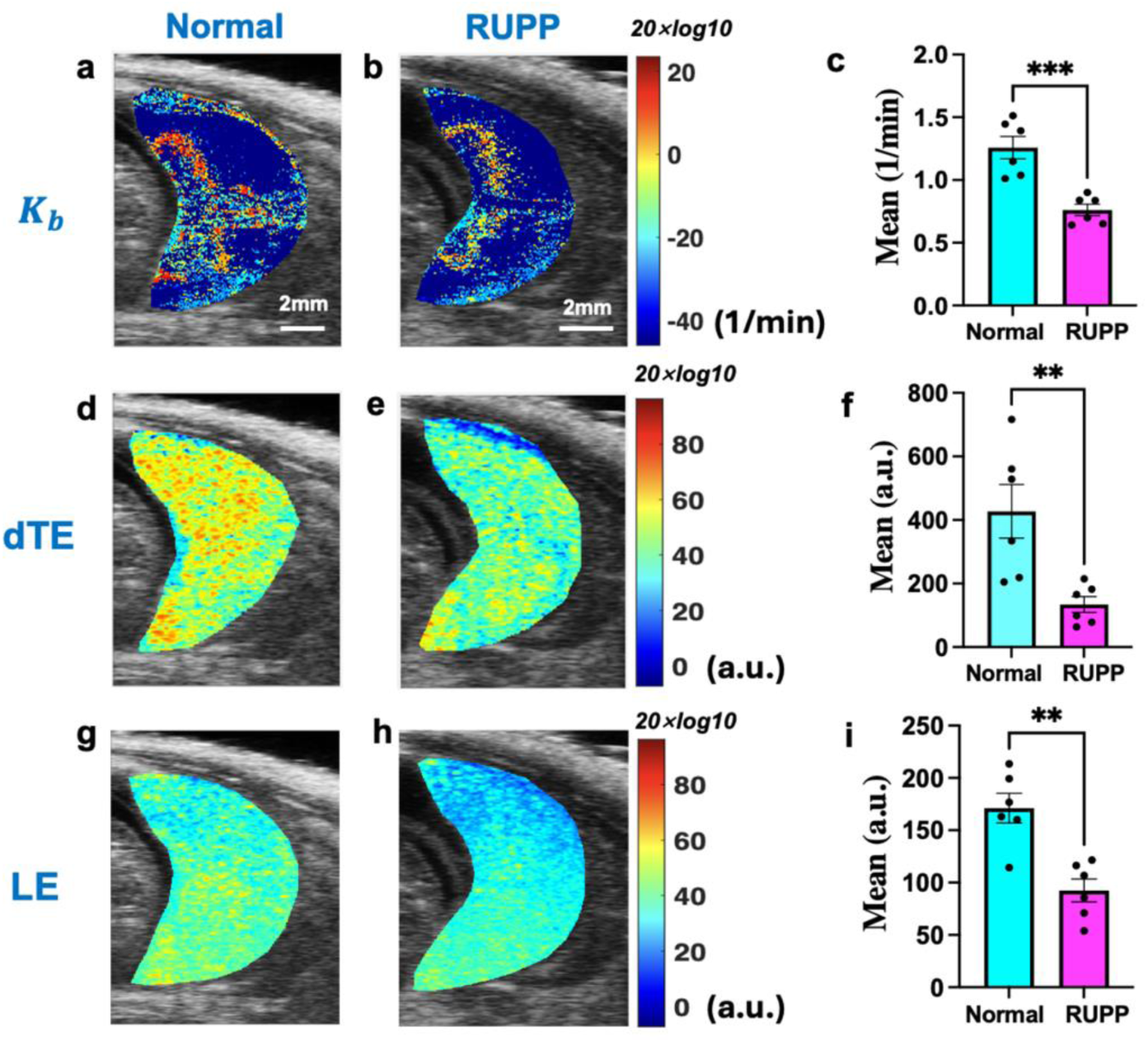
Parametric maps of bound MBs. **(a, b)** *K_b_***, (d, e)** dTE, (**g, h)** LE. The parameter values shown in all parametric maps are obtained through a 20*log10 transformation. NP **(a, d, g)** vs RUPP **(b, e, h)**. All methods show higher binding MB values in NP placentas. **(c, f, i)** The histograms compare mean values in three methods: NP (n=6) significantly higher than RUPP (n=6). Each point in **(c)** shows the mean value of top half of each placenta, where signals above noise were recorded within the first 2 mins post-injection. Each point in **(f, i)** represents the mean value of the entire placenta region for each rat. An unpaired two-sided t-test indicates statistically significant differences, denoted as (**p< 0.01, ***p< 0.001). The statistical power was analyzed using G*Power, and for *K_b_*, α > 0.95, for dTE, α > 0.85, and for LE, α > 0.9.

Figure 4a illustrates *K_b_*, dTE, and LE values for a representative NP placenta and a representative RUPP placenta. The histogram displays the pixel distribution of the derived parameters. The histogram of *K_b_* presents three peaks: the highest at −50 (no bound MBs), and two others corresponding to central and labyrinth zones (more attached MBs). LE and dTE histograms display single peaks, offering less insight into placental molecular heterogeneity.

**Figure 4.**
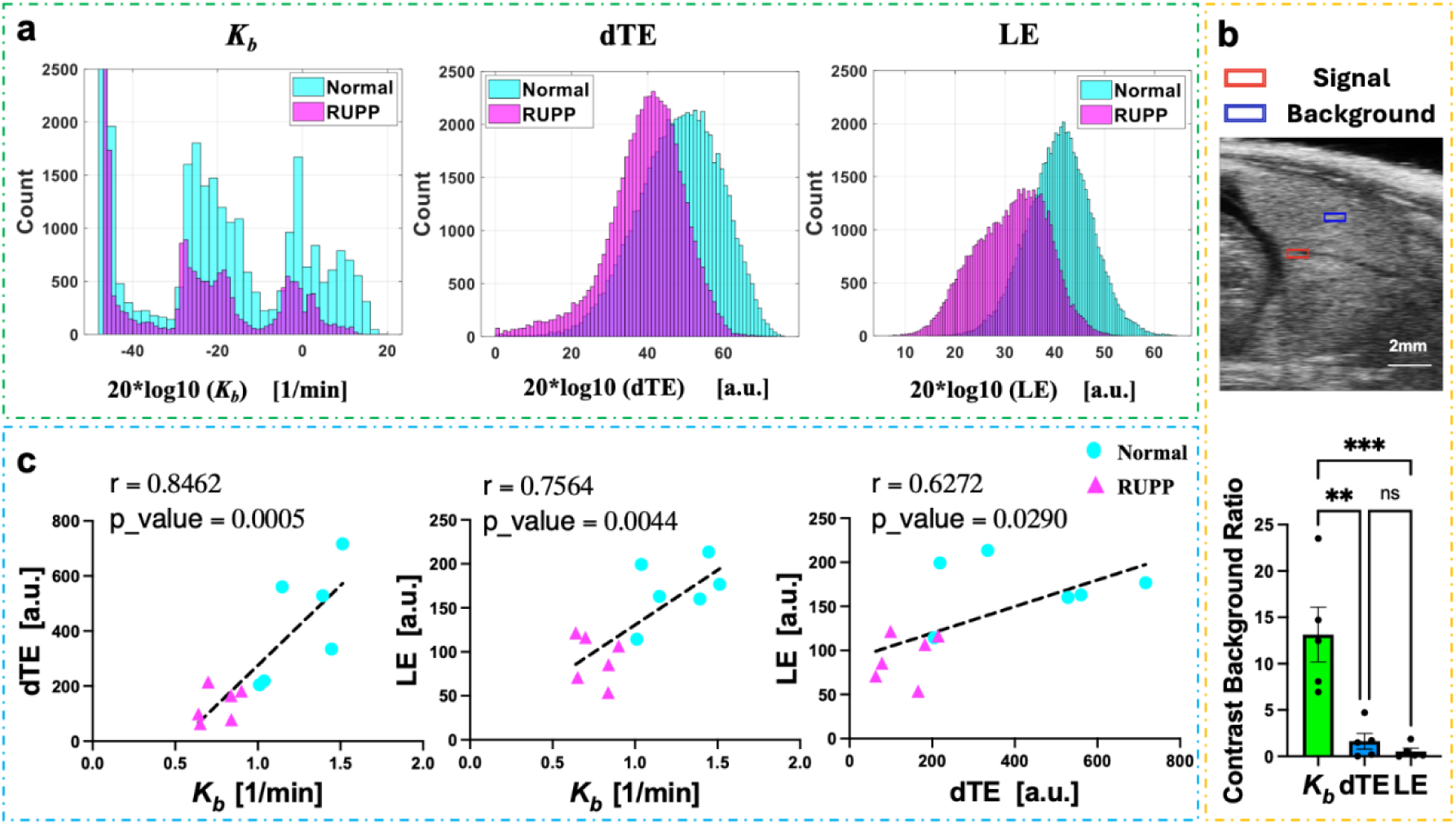
**(a)** Histograms of the *K_b_*, dTE, and LE parameters in a normal (blue) and RUPP (purple) placenta. **(b)** Contrast-to-background ratio comparison (means ± SEM, n=5 normal pregnant rats; **p< 0.01, ***p< 0.001). **(c)** Regression and correlation of the three methods, where r is the correlation coefficient. Scatter plots show the relationships of dTE and LE versus *K_b_*. Each point represents the mean value of each placenta ROI, calculated as described for Figure 3c, f, and i. The regression between dTE and *K_b_* has the highest significance.

### Comparison of the Contrast-to-Background Ratio Between BCM, dTE, and LE Methods

Figure 4b, a 10x10 pixel region (red square) was positioned within the central arterial canal of the placenta. The central arterial canal, the largest maternal vessel in the rat placenta, exhibited the most rapid signal growth, reaching its maximum value faster than other areas. Another 10x10 pixel region (blue square) was chosen as the background, located in an area with fewer blood vessels, where signal growth was significantly slower. The contrast-to-background ratios of the *K_b_*, dTE, and LE parameter maps were compared for all normal rat placentas. The *K_b_* parameters had higher average contrast-to-background ratios, indicating better vascular structure distinction. A one-way ANOVA was used to compare the ratios between the three methods. Results showed significant differences between *K_b_* and LE. In Figure 4b, the histogram plots the placental parameter averages of five normal rats (n=5). Six placentas were imaged, with one placenta excluded from further analysis due to the absence of a visible hypoechoic central arterial canal on the B-mode image, preventing an accurate segmentation.

Figure 4c illustrates the correlation and regression results for three comparisons: *K_b_* vs. dTE (\**p* < 0.05), *K_b_* vs. LE (\**p* < 0.05), and dTE vs. LE (\**p* < 0.05). The correlation and regression analyses (performed in GraphPad Prism Version 10.2.0, GraphPad, San Diego, CA), revealed significant correlations between the three methods.

### Expression Levels of α_ν_β_3_ Changed in Placentas of RUPP Animals

The IHC staining for α_ν_β_3_, highlighting vascular endothelial cells, in NP and RUPP placentas is shown in Figures 5a and 5b, respectively. In Figure 5c, each point in the histogram represents the average value of α_ν_β_3_ integrin protein expression from all slices for each rat. RUPP placentas exhibited a significant reduction of the staining area fraction (Figure 5c), and downregulated α_ν_β_3_ mRNA expression (Figure 5d) compared to NP. This finding confirms the targeted CEUS imaging measures impaired vascular development and remodeling in the placenta.

**Figure 5.**
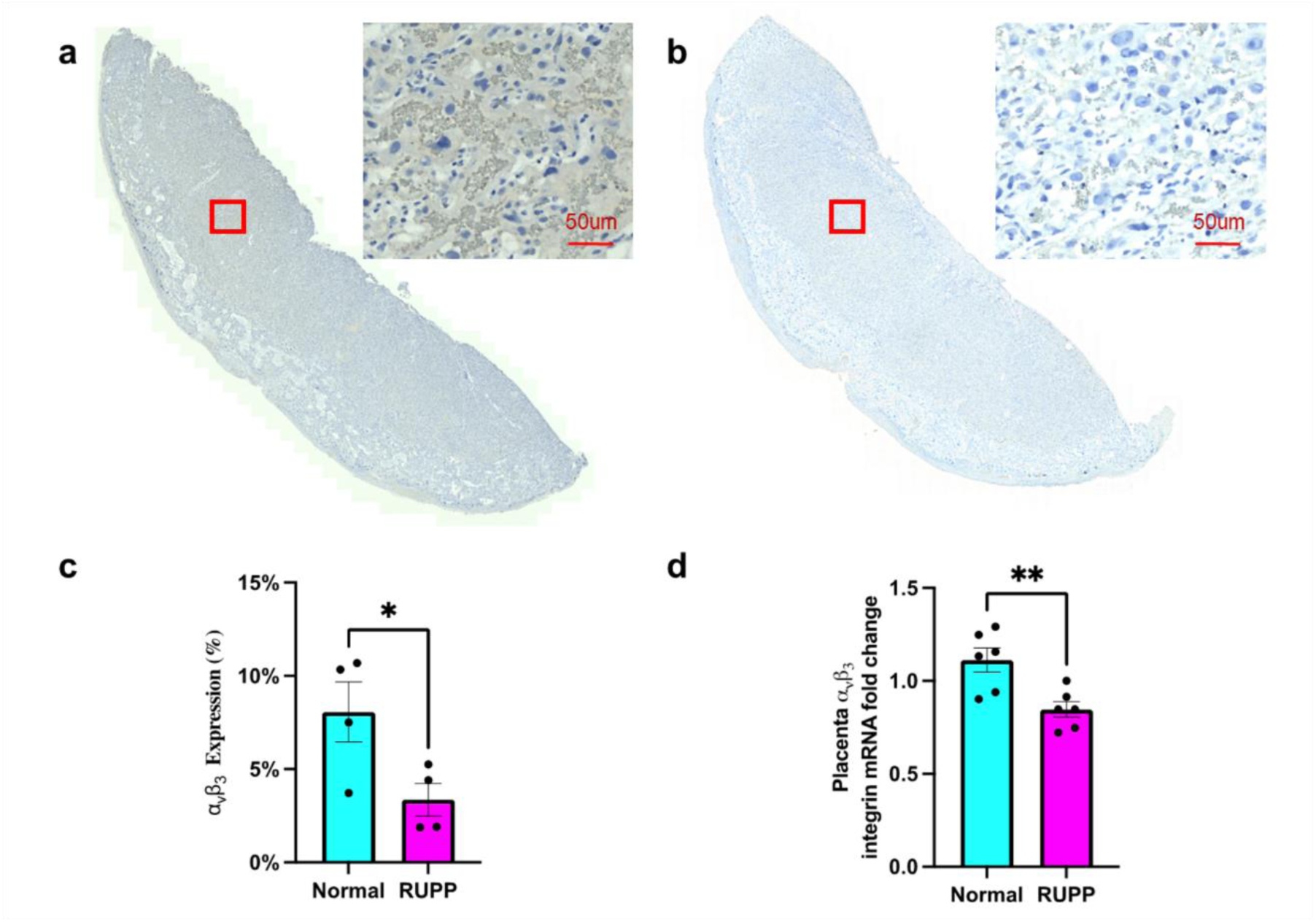
α_ν_β_3_ integrin expression in NP vs RUPP placentas. **(a)** IHC staining in NP placenta. **(b)** IHC staining in RUPP placenta, showing decreased intensity. **(c)** Quantification of α_ν_β_3_ integrin protein expression. Each data point represents the mean of 3-4 slices from a single placenta from each rat subject. (n=4 rats; mean ± SEM; *p< 0.05). **(d)** Relative α_ν_β_3_ mRNA levels (n=6 each; normalized to β-Actin; mean ± SEM; **p<0.01).

## Discussion

Evaluating abnormal placental development *in vivo* is crucial for identifying pregnancy-related complications like PE. During PE, insufficient remodeling of the spiral arteries leads to incomplete dilation of the capillaries, narrowing the blood vessel walls and inhibiting placental blood flow [35, 36]. This inadequate blood supply reduces oxygen and nutrient delivery to the placenta, adversely affecting vessel architecture growth [37, 38]. To address this, we used T-CEUS imaging as a non-invasive technique to analyze *in vivo* placental vascular development.

Our results show a reduced binding constant *K_b_* in PE, supporting our hypothesis that the BCM model provides a more precise analysis of vascular changes in placental tissue. By fitting each pixel’s TIC into the mathematical model, we obtained a detailed mapping of angiogenesis and MB binding parameters. The decreased *K_b_* in PE indicates reduced angiogenesis in the placenta [39], aligning with previous studies linking PE to impaired placental development [40, 41]. Specifically, the reduced MB attachment to α_ν_β_3_ integrin in RUPP animals suggests significant impairment in placental vascular development, potentially contributing to adverse outcomes in PE. Furthermore, our findings highlight the potential of the BCM model to enhance the diagnosis and monitoring of pregnancy-related complications through detailed, quantitative assessment of placental vascular health, potentially enabling early detection and intervention in PE.

Placental circulation begins from the maternal uterine artery through the radial artery, to the spiral artery [42], forming the central arterial canals within the placenta, which perfuse the labyrinth zone [43]. The labyrinth zone is the primary location of maternal-fetal exchange [43] and exhibits high cellular proliferation [44]. Our study observed a higher TCA concentration in the central canal and labyrinth zone in NP compared to the RUPP model, quantitatively linking PE to altered placental structure. The decreased binding constant *K_b_* and reduced α_ν_β_3_ integrin attachment in the PE model indicate diminished angiogenesis. This aligns with previous research, showing abnormal placental development in PE, characterized by decreased blood vessels, reduced vessel remodeling, and increased apoptosis in the labyrinth zone [45]. The impaired angiogenesis in the labyrinth zone suggests that adverse maternal and fetal outcomes associated with PE are due to the compromised exchange of gases, nutrients, and waste products.

By providing a spatially quantitative assessment of these changes, our study shows the potential of using targeted CEUS imaging and BCM analysis to enhance the understanding, diagnosis, and monitoring of PE and other pregnancy-related complications. More conventionally used late enhancement (LE) and differential targeted enhancement (dTE) methods were compared to the BCM results. The LE method requires a 10-minute data acquisition time to allow the freely circulating microbubble to clear, and the contrast signals are dominated by the attached microbubble. LE enables qualitative molecular imaging; however, the targeted microbubble signals are not distinct from the freely circulating microbubbles; it is not appropriate for quantitative contrast imaging of the placenta [13]. dTE methods require an extended acquisition time and a burst pulse, potentially causing bioeffects. Additionally, both semi-quantitative methods only analyze several frames at corresponding time points to analyze the data, which reduces spatial heterogeneity when compared to the bicompartmental model.

Unlike the two traditional methods that select specific time points, the BCM fits the time-intensity curve (TIC) of each pixel over a continuous time range. This provides a more precise fitting method and better captures the dynamic changes in the particularly heterogenous vasculature of the placenta. The BCM method differentiated a wider range of placental anatomical structures. The higher contrast-to-background ratio enables the distinction of more intricate internal vascular structures, thereby providing a wealth of additional anatomical information, as illustrated in Figure 4. Regression and correlation analyses of the three methods are shown in Supplementary Figure 6, demonstrating that, at least with this limited sample set, *K_b_* uniquely differentiates between normal and RUPP cases, while dTE and LE do not provide separation.

There are limitations of our studies. In calculating the attached MBs, we cannot avoid the possibility of secondary binding of microbubbles. This can also affect our calculation of the number of bound MBs. However, our purpose is to distinguish between Normal and RUPP groups, so although it may affect the specific numerical values of MB binding, it does not impact our ability to monitor preeclampsia. This preclinical study used an ultrasound contrast agent which is not currently clinically approved. There is however a VEGFR2-targeted microbubble, BR55, which is commercially available and in clinical trials [21, 46]; we anticipate additional targets being clinically available in the future, allowing for translation of these methods towards the clinic. However, for future clinical applications, additional preclinical testing and validation studies would be needed.

CEUS is used clinically to diagnose atherosclerosis, aortic aneurysms, and thrombosis. Agents are typically expelled from the body through respiration [47]. Clinically approved microbubbles have good safety profiles and low rates of adverse reactions [48]. Although CEUS is not currently FDA-approved for clinical use in pregnant women, pregnant women have been exposed to microbubbles during diagnostic imaging for other conditions, and no contrast agent was detectable in the fetus [49]. In studies conducted on rats, injection of SonoVue™ microbubbles into the rats did not enhance the passage of large molecules across the placental barrier to reach the fetus [50–52].

Various imaging techniques have been used to evaluate PE. MRI-based methods, such as placental diffusion-derived vascular density (DDVD) analysis [53], and blood oxygen level-dependent (BOLD) MRI can detect placental dysfunction and ischemia [54], but their cost and long imaging times limit clinical use. Photoacoustic imaging measures the placental oxygen saturation to detect placental ischemia and determine the presence of fetal growth restriction [55], but currently has limited availability and limited imaging depth. Doppler ultrasound can be used to monitor blood flow in the uterine artery and umbilical cord but lacks sensitivity for intraplacental flow. The ability to non-invasively monitor these changes *in vivo* would be a significant advancement in the management of PE and other related conditions, ultimately improving maternal and fetal health outcomes.

## Conclusions

Our study highlights the potential of molecular contrast-enhanced ultrasound imaging, accompanied by bicompartmental modeling, for the diagnosis of placental dysfunction-related diseases. We have demonstrated quantitative *in vivo* measurement of angiogenesis-related molecular expression within the placenta of a rat model of preeclampsia and demonstrated that the quantified reduction in molecular binding is consistent with the reduced placental α_ν_β_3_ integrin expression. The demonstrated techniques could be applied to analyze *in vivo* placental molecular expression in other pregnancy-related diseases, such as fetal anemia, hypertension, and gestational diabetes.

## Author Contributions

Lili Shi, Dylan Lawrence, and Carolyn L. Bayer designed the study; Lili Shi, Allan K. N. Alencar and Kenneth F. Swan performed the image acquisitions; Lili Shi performed the image analysis; Lili Shi interpreted the results and wrote the initial manuscript; all authors contributed to and approved the final manuscript.

## Acknowledgements

We would like to express our sincere appreciation to Ashli Olson of the Pathology Core Laboratory at the Tulane University Health Science Center for her assistance with the immunohistology, as well as the Tulane University Brain Institute Cell and Tissue Imaging Core for access to Zeiss Axio Scan.Z1 Slide Scanner.

## Ethical Approval

This study followed all applicable institutional and/or national guidelines for the care and use of animals.

## Data Availability

All research data and computer codes are available from the corresponding author upon request.

