## Supplementary Figure 1, 2, 3, 4, 5, 6 for "Quantifying Molecular Changes in the Preeclamptic Rat Placenta with Targeted Contrast-Enhanced Ultrasound Imaging"

**Journal: Molecular Imaging and Biology**

Lili Shi,[1] Allan K. N. Alencar,[1] Kenneth F. Swan,[2]
Dylan J. Lawrence,[1] Gabriella Pridjian,[2] Carolyn L. Bayer,[1]

1. Department of Biomedical Engineering, Tulane University, New Orleans, LA, 70118, USA.
2. Department of Obstetrics & Gynecology, Tulane University, New Orleans, LA, 70112, USA.

**Corresponding author:**

Carolyn L. Bayer

Department of Biomedical Engineering,

Tulane University,

500 Lindy Boggs Center,

New Orleans, LA, 70118, USA


### Supplemental Materials and Methods

*Reduced Uterine Perfusion (RUPP) Model of Preeclampsia*

The Institutional Animal Care and Use Committee at Tulane University approved all animal studies, which were conducted following the guidelines outlined in the NIH Guide for the Care and Use of Laboratory Animals. Timed-pregnant Sprague Dawley rats, aged approximately 8 weeks and weighing around 250 g, were purchased from a commercial vendor (Charles River Laboratories, Boston, MA). Animals were received on gestational day (GD) 11 and housed individually in a controlled environment with a temperature of 21°C, maintained on a 12-hour light/dark cycle, and provided access to food and water ad libitum. On GD14, animals were randomly divided into two cohorts: Normal pregnant (NP; n = 6), and RUPP (n = 6). The RUPP procedure was performed under anesthesia, with 2-3% isoflurane administered through inhalation, and involved placing silver clips on the lower abdominal aorta and branches of the ovarian arteries to induce placental ischemia as previously published (Supplementary Figure 1) [27–29]. Perioperative pain was managed by administering carprofen (Rimadyl, injectable, 5 mg/kg, Zoetis, Parsippany, NJ) via subcutaneous injection, while postoperative pain was controlled daily through oral administration of carprofen (Rodent MD's, Rimadyl, 2 mg/tablet, Bio-Serve, Flemington, NJ). Rats were monitored once a day after surgery until GD18. Animals were euthanized by decapitation with prior isoflurane anesthesia at the completion of all experiments.

*Preparation of Targeted Contrast Agent (TCA)*

Vevo Targeted-Ready Micro Markers (Fujifilm, VisualSonics) were utilized for this experiment. These MBs can function as non-targeted contrast agents or be conjugated with peptides to target specific biomolecules on the surface of endothelial cells. Following the manufacturer's protocol, 0.7 mL of saline was added to the Micro Marker vial and gently shaken for one minute to ensure thorough reconstitution. Subsequently, 20 µg of biotin-Arginine-Glycine-Aspartate-OH TFA salt (RGD) peptide was diluted in 300 µl of saline. The entire peptide solution was then injected into the Vevo Micro Marker vial. The resulting TCA are designed to bind to $\alpha_v\beta_3$ integrin [30].

*Gene expression*

After euthanasia, placentas were collected and weighed before being frozen in liquid nitrogen and stored at -80 °C. Real-time quantitative polymerase chain reaction (RT-qPCR) was employed to assess mRNA expression levels in the placental tissues. Specifically, the mRNA levels of $\alpha_v\beta_3$ integrin and β-actin were quantified. The extraction of mRNA from placental tissues (30 mg) was conducted utilizing the TRIzol Reagent (Ambion-Life Technologies, USA) at a ratio of 1 mL TRIzol Reagent per sample. Tissue disruption was facilitated by a Fisher Scientific bead beater, followed by purification using the RNeasy mRNA extraction kit (Qiagen, Germany). The concentration of mRNA was determined using the Nanodrop 1000 Spectrophotometer. Subsequently, 10 μg of mRNA underwent DNase treatment with the Turbo-DNA-free kit (Invitrogen, Thermo Fisher Scientific, USA). Following this, 1 μg of mRNA was reverse-transcribed into cDNA using the iScript cDNA Synthesis Kit (Bio-Rad, USA) in accordance with the manufacturer's instructions. SsoAdvanced Universal SYBR Green Supermix (Bio-Rad, USA) was used for the RT-PCR reaction. RT-qPCR was performed using the CFX Connect Real-Time System (Bio-Rad, USA). The PCR protocol consisted of an initial denaturation step at 95°C for 3 minutes, followed by 40 cycles of a two-step amplification process: denaturation at 95°C for 10 seconds and annealing at a temperature specific to each primer for 30 seconds. For each experiment, melting curve analysis was conducted to validate the specificity of the PCR products. β-Actin was used as the housekeeping gene for the normalization of $\alpha_v\beta_3$ integrin mRNA expression. The fold change for $\alpha_v\beta_3$ integrin mRNA was calculated using the $2^{\wedge(-\Delta\Delta CT)}$ method of analysis.

*Immunohistochemistry*

Post-imaging on GD18, animals were euthanized by decapitation under isoflurane anesthesia, and one placenta from each animal was collected for immunohistochemistry. The placentas were fixed in formalin, embedded in paraffin blocks, and cut into 4 μm sections. Immunohistochemical staining was performed using the Horseradish Peroxidase method (HRP) on placental sections closest to the placental midpoint – i.e. approximately the same region of CEUS image acquisition. Placental sections were incubated with mouse Polyclonal $\alpha_v\beta_3$ integrin (1:200 dilution, Bioss antibodies, bs-1310R) and secondary HRP-polymer (Rabbit-On-Rodent HRP-polymer, Biocare Medical, Pacheco, CA). Slides were then incubated with 3,3′-diaminobenzidine (DAB) for 5-min and rinsed in a running water bath, and counterstained with hematoxylin.

Slides were scanned using a Zeiss Axio Scan.Z1 Slide Scanner (Carl Zeiss AG, Oberkochen, Germany) and images were exported to Fiji ImageJ analysis software for processing. The placental border was first manually segmented from the $\alpha_v\beta_3$ stained images, and a color deconvolution was applied to separate the DAB and hematoxylin signals. The expression of $\alpha_v\beta_3$ integrin was quantified as the percent area fraction of DAB staining compared to the total area of the placental section. All histological image analysis was performed by users blinded to the study.

### Supplemental Results

*Quantitative analysis of targeted contrast agent (TCA).*

Supplementary Figures 2, 3, and 4 show the parametric map of $K_b$, dTE, and LE values for all placentas. Compared to the Normal group (Supplementary Figure 2a, b, c, d, e), the number of attached MBs in RUPP placentas (Supplementary Figure 2f, g, h, i, j) significantly decreased, while normal placentas showed higher attachment. The semi-quantitative methods (Supplementary Figures 3 and 4) also exhibited higher values in NP placentas. Supplementary Figure 5 shows a conventional dTE (dTEc) which calculates the signal difference just before and just after the burst. The dTEc provides reduced contrast in comparison to the dTE methods used in the main manuscript. Supplementary Figure 6 shows ROC curves of three parameters, indicating that $K_b$ provides the best distinction between NP and RUPP.

### Supplemental Figures

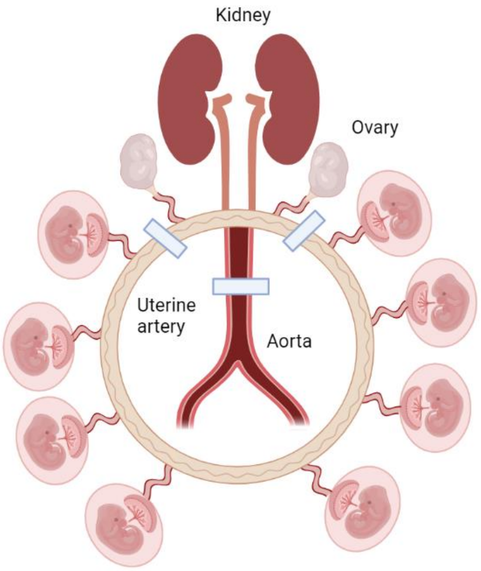


**Supplementary Figure 1.** Schematic diagram of the Reduced Uterine Perfusion Pressure (RUPP) model. On gestational day (GD) 14, the lower part of the abdominal area is opened. A silver clip (d=0.203 mm) is carefully placed on the central aorta, and two silver clips (d=0.1 mm) are placed on the uterine arteries. This procedure limits the blood supply to the uterus and mimics the symptoms of preeclampsia.

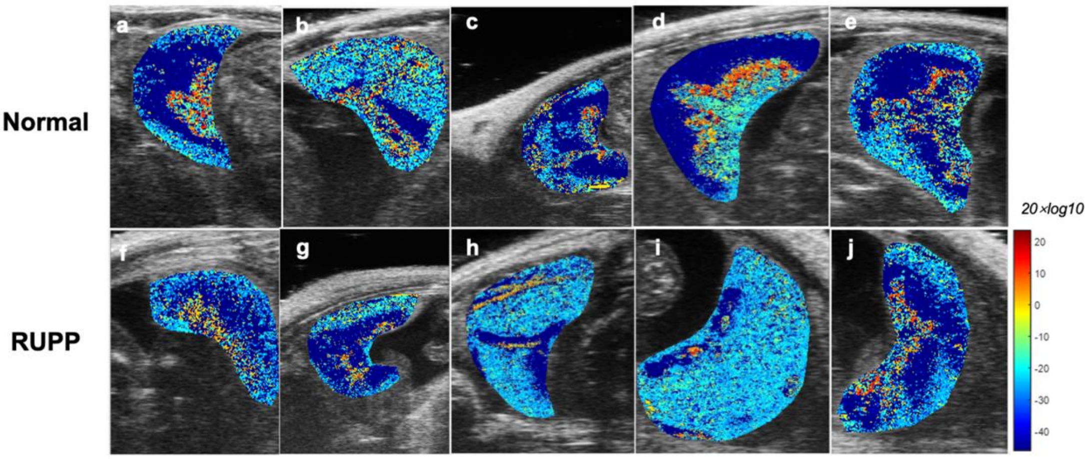


**Supplementary Figure 2.** Parametric maps of BCM methods. **(a, b, c, d, e)** Normal placenta $K_b$ maps and **(f, g, h, i, j)** RUPP placenta $K_b$ maps. Compared to normal rats, the RUPP group shows decreased $K_b$ values, suggesting lower bound microbubbles.

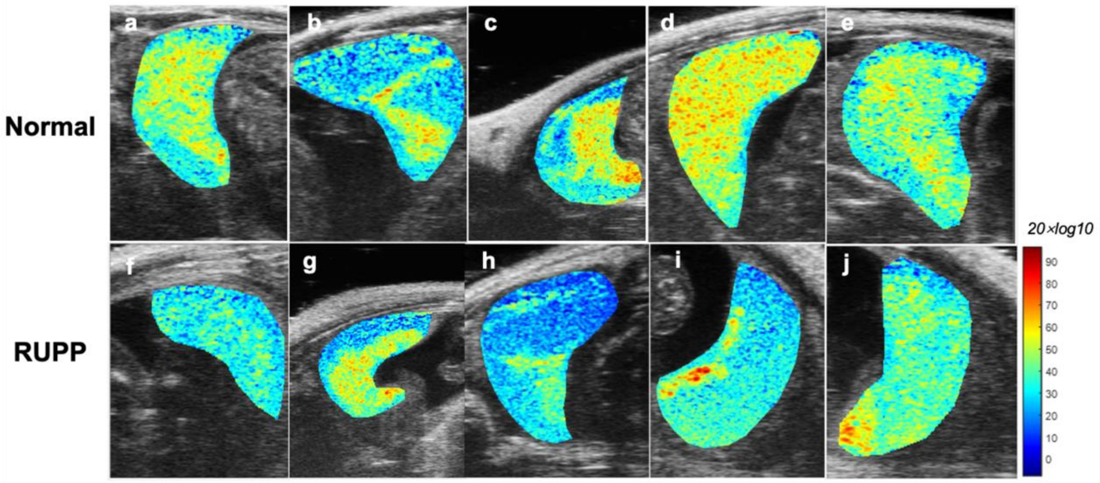


**Supplementary Figure 3.** Parametric maps of dTE methods. **(a, b, c, d, e)** Normal placenta dTE maps, and **(f, g, h, i, j)** RUPP placenta dTE maps. Compared to normal rats, the RUPP group shows decreased values, suggesting lower bound microbubbles.

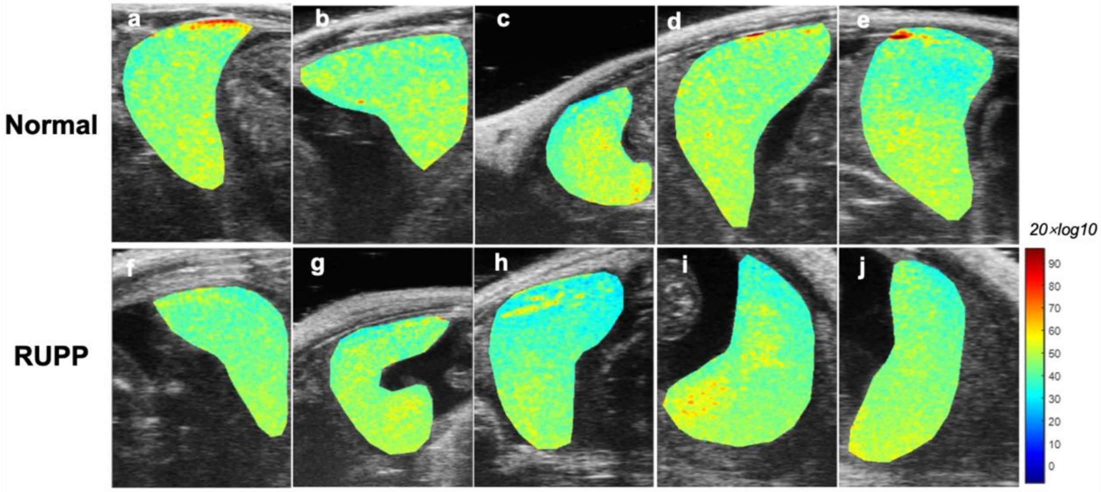


**Supplementary Figure 4.** Parametric maps of LE methods. **(a, b, c, d, e)** Normal placenta LE maps, and **(f, g, h, i, j)** RUPP placenta LE maps. Compared to normal rats, the RUPP group shows decreased values, suggesting lower bound microbubbles.

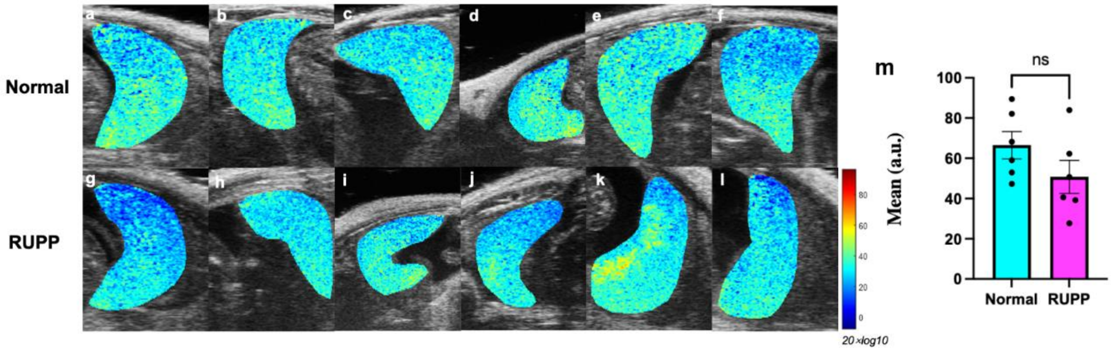


**Supplementary Figure 5.** Parametric maps of dTEc. **(a, b, c, d, e, f)** Normal placenta, and **(g, h, i, j, k, l)** RUPP placenta dTEc maps. **(m)** Bar chart of the mean and standard deviation of the dTEc for the entire placenta region of each rat. Statistical analysis used an unpaired two-sided t-test; dTEc was not significantly different for normal and RUPP placentas.

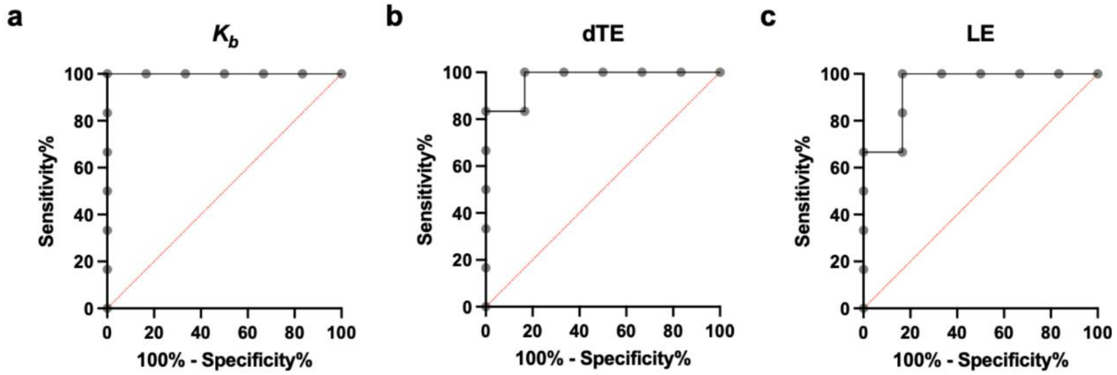


**Supplementary Figure 6.** Receiver operating characteristic (ROC) curves for the (a) $K_b$ , (b) dTE, and (c) LE parameters (NP, n=6; RUPP, n=6). The red line indicates random guessing.

Comparing the ROC curves of the three methods, $K_b$ performs better than dTE, which in turn outperforms LE.
